# Dysregulated nuclear export of the late herpes simplex virus 1 transcriptome through the vhs-VP22 axis uncouples virus cytopathic effect and virus production

**DOI:** 10.1101/2022.11.02.514834

**Authors:** Kathleen Pheasant, Dana Perry, Emma Wise, Vivian Cheng, Gillian Elliott

**Affiliations:** Section of Virology, Department of Microbial Sciences, University of Surrey, Guildford, United Kingdom

## Abstract

Herpes simplex virus 1 (HSV1) expresses its genes in a classical cascade culminating in the production of large amounts of structural proteins to facilitate virus assembly. HSV1 lacking the virus protein VP22 (Δ22) exhibits late translational shutoff, a phenotype that has been attributed to the unrestrained activity of the virion host shutoff (vhs) protein, a virus-encoded endoribonuclease which induces mRNA degradation during infection. We have previously shown that vhs is also involved in regulating the nuclear-cytoplasmic compartmentalisation of the virus transcriptome, and in the absence of VP22 many virus transcripts are sequestered in the nucleus late in infection. Here we show that despite expressing minimal amounts of structural proteins and failing to plaque on human fibroblasts, the strain 17 Δ22 virus replicates and spreads as efficiently as Wt virus, but without causing cytopathic effect (CPE). Nonetheless, CPE-causing virus spontaneously appeared on Δ22-infected human fibroblasts, and four viruses isolated in this way had all acquired point mutations in vhs which rescued viral mRNA export and late protein translation. However, unlike a virus deleted for vhs, these viruses still induced the degradation of cellular mRNA, suggesting that vhs mutation in the absence of VP22 is necessary to overcome a disturbance in mRNA export rather than mRNA degradation. The ultimate outcome of secondary mutations in vhs is therefore the rescue of virus-induced CPE caused by late protein synthesis, and while there is a clear selective pressure on HSV1 to mutate vhs for optimal production of late structural proteins, the purpose of this is over and above that of virus production.

**Author Summary:** HSV is a human pathogen that lytically infects cells of the epidermis. Following viral genome replication, structural proteins are produced in abundance to enable the rapid assembly and release of large quantities of infectious progeny. Infected cells also exhibit cytopathic effect (CPE), morphological changes that are exemplified by cell rounding and the breakage of cell-to-cell contacts, facilitating virus dissemination. Here we show that HSV1 with a mutation that results in the nuclear retention of viral mRNA and concomitant shutdown of late protein synthesis, also fails to cause CPE. However, unexpectedly, we found that this virus is still able to release large numbers of infectious virus which can spread between cells without any evidence of cell damage. Nonetheless, despite efficient virus productivity, this virus spontaneously mutates to rescue late protein production and CPE, with mutations mapping to the process of mRNA export. There is therefore a clear selective pressure on HSV1 to optimize the synthesis of late structural proteins, but the purpose of this is over and above that of virus production, a result that has implications for why viruses in general express such large amounts of structural proteins.

## Introduction

Herpes simplex virus type 1(HSV1) expresses its genes in a classical cascade of gene expression during lytic infection, comprising immediate-early, early and late genes [1]. In general, the late genes encode virus structural proteins and are transcribed predominantly from replicated DNA genomes, leading to a large burst of late protein synthesis for optimal virus assembly. Deletion of the HSV1 UL49 gene which encodes the tegument protein VP22 [2] results in a virus that exhibits translational shutoff of late protein synthesis [3, 4], and in many systems is detrimental to virus propagation [4–6]. This translational shutoff is not a consequence of enhanced host responses such as the stress response kinase protein kinase R. Rather, it correlates with the failure of late viral transcripts to be exported from the nucleus of cells infected with the Δ22 virus, as demonstrated in our previous studies using mRNA FISH, thereby preventing their translation in the cytoplasm [4].

A clue to the mechanism by which viral transcripts are retained in the nucleus of Δ22-infected cells came from the observation that spontaneous secondary mutations frequently arise in the UL41 gene of the Δ22 genome [4, 6, 7], a gene which encodes the virion host shutoff (vhs) protein [8]. These mutations rescue the deleterious effect of VP22 deletion on late protein translation, restoring plaque formation [4, 6, 7]. The vhs protein is an endoribonuclease which induces the degradation of cellular mRNA during HSV1 infection through its endoribonuclease cleavage of cytoplasmic mRNAs followed by Xrn1 exonuclease degradation [9], and regulates the transition from IE to E and L gene expression [10, 11]. It was therefore originally proposed that VP22 is required to quench vhs-induced mRNA degradation at later times in infection and that in the absence of VP22, vhs endoribonuclease activity is lethal [6]. Nonetheless in our hands, infection of human fibroblasts with a Δ22 virus did not result in unrestrained mRNA degradation compared to Wt infection [4]. Moreover, in that study we also demonstrated that in Wt infection, IE and E transcripts were retained in the nucleus at late times, but in cells infected with a Δvhs virus all classes of transcripts were exported to the cytoplasm [4], suggesting that the vhs endoribonuclease is involved in regulating mRNA export, and providing a link between mRNA degradation in the cytoplasm and mRNA retention in the nucleus. The relative compartmentalisation of the virus transcriptome was also mirrored by the localisation of the polyA binding protein PABPC1, a protein that has a steady-state cytoplasmic localisation but shuttles between the cytoplasm and nucleus to bind polyadenylated mRNAs ready for export [12, 13]. Once mRNA in the cytoplasm has been turned over, PABPC1 recycles back to the nucleus, and in the presence of functional vhs, PABPC1 accumulates there in an endoribonuclease dependent fashion [14, 15].

Here we report the unexpected result that despite translational shutoff and lack of plaque formation in primary human fibroblasts, HSV1 lacking VP22 replicates, spreads and produces as much infectious progeny virus as Wt virus without causing any cytopathic effect (CPE) in these cells. Nonetheless, CPE-inducing virus rapidly appeared as plaques in Δ22-infected fibroblasts and these rescued viruses had all acquired mutations in vhs which restored nuclear export of viral mRNA during infection rather than abrogating mRNA degradation. This suggests that despite efficient virus propagation in the absence of VP22, there is pressure on the virus to mutate vhs and restore late protein translation and concomitant CPE, over and above what is required for virus production. These results have implications for understanding why this and potentially other viruses express such large amounts of late virus proteins.

## Results

### HSV1 lacking VP22 replicates and spreads in primary human fibroblast cells without causing cytopathic effect

Deletion of the VP22-encoding gene (UL49) has been shown to be detrimental to HSV1, resulting in extreme late translational shutoff [4, 6, 7]. Our own Δ22 virus based on strain 17 fails to plaque on primary human fibroblasts (HFFF) as late as 5 dpi (Fig 1A). The efficiency of viral DNA replication in Δ22 infection was measured by harvesting at 2 or 16 h after infection and determining the relative viral DNA copy number by qPCR of the virus gene *UL48*, to reflect input viral DNA (2 h) or viral DNA replication (16 h). Although there was less input DNA in Δ22 infected cells, the relative increase in genome copies was similar in Wt and Δ22 infected cells at 16 h, indicating that the absence of VP22 has little effect on genome replication (Fig 1B), and that the block to virus production occurs at a later stage. Western blotting of HFFF cells infected at high multiplicity confirmed that a range of virus envelope proteins are poorly expressed (Fig 1C), in line with the previously demonstrated translational shutoff in these cells [4], providing an obvious explanation for the inability of this virus to form plaques in HFFF cells. Immunofluorescence of infected cells revealed that the IE protein ICP4, which localises in a distinctive cytoplasmic punctate pattern late in Wt infection, was restricted to the nucleus in Δ22 infected cells (Fig 1D, ICP4) while the envelope protein glycoprotein E (gE) was concentrated in a juxtanuclear compartment rather than progressing to the plasma membrane as it does in Wt infection (Fig 1D, gE). These results suggest there is a block to late protein trafficking in the absence of VP22. However, despite these obvious defects in Δ22 infection, we found no significant difference in the growth kinetics of Δ22 compared to Wt virus in HFFF in a one-step growth curve (Fig 1E). Moreover, low magnification imaging of HFFF cells infected with Δ22 (which expresses GFP in place of VP22) showed that while all cells were GFP positive after 20 h, there was no sign of the classical HSV1-induced cytopathic effect (CPE) of cell-rounding, which was evident in cells infected with Wt or HSV1 expressing GFP fused to VP22 (GFP-22) infected cells (Fig 1F). By contrast, HSV1 expressing GFP in place of UL34, a protein essential for nuclear egress [16], exhibited CPE similar to Wt and GFP-22 (Fig 1F), indicating that even though this virus is unable to export capsids to the cytoplasm or assemble progeny virions, it is still able to cause CPE.

**Fig. 1.**
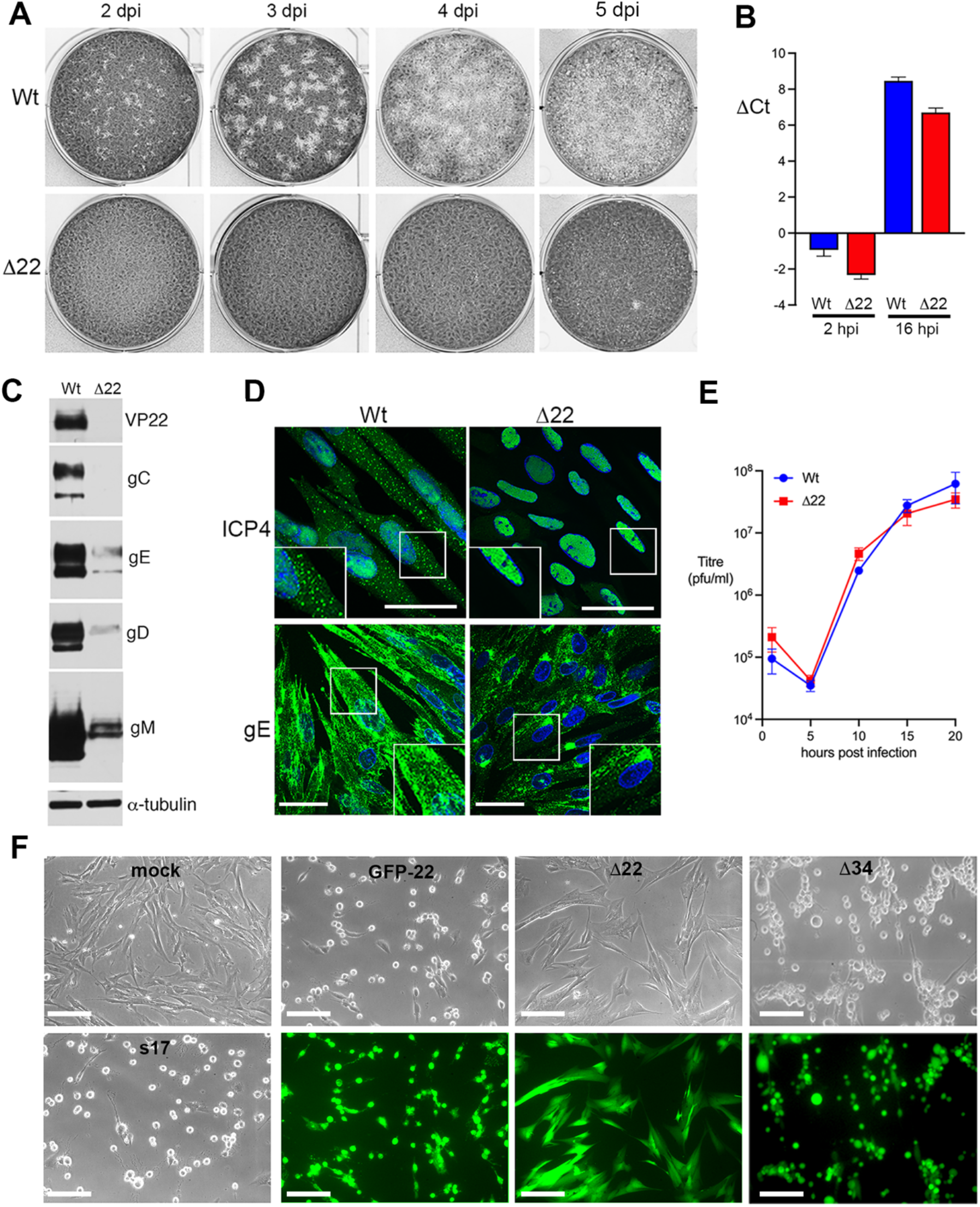
HSV1 replicates in primary human fibroblasts in the absence of VP22 without causing CPE. **(A)** HFFF cells were infected with approximately 50 pfu of Wt s17 and Δ22 viruses, fixed at 2, 3, 4 and 5 days (dpi) and stained with crystal violet. **(B)** HFFF cells were infected with Wt s17 or Δ22 virus at a multiplicity of 3, acid washed at 1 hpi, then DNA was isolated at 2 or 16 hpi. qPCR was performed for gene *UL48* to determine the relative virus DNA copy number represented as ΔCt to 18s rDNA (mean±SEM, *n* = 3). **(C)** HFFF cells infected with Wt (s17) or Δ22 viruses at MOI 2 were harvested at 16 hpi and analysed by SDS-PAGE and Western blotting with antibodies as indicated. **(D)** HFFF cells infected with Wt (s17) or Δ22 viruses at MOI 2 were fixed at 16 hpi and analysed by immunofluorescence with antibodies to the IE protein ICP4 and the L protein glycoprotein E (gE), both in green. Nuclei were stained with DAPI (blue). Scale bar = 50 μm. **(E)** HFFF cells were infected with Wt s17 or Δ22 virus at a multiplicity of 2, total virus harvested every 5 h up to 20 h and titrated onto Vero cells (mean±SEM, *n* = 3). **(F)** HFFF cells grown in 6-well plates were left uninfected (mock) or infected at a multiplicity of 2 with Wt s17, HSV1 GFP-22, Δ22 expressing GFP or Δ34 expressing GFP. After 20 h the cells were imaged live using brightfield and fluorecence where appropriate. Scale bar = 100 μm.

Given that the Δ22 virus does not plaque on HFFF, we next investigated its ability to spread in these cells. A multi-step growth curve was carried out by infecting HFFF cells at a multiplicity of 0.01 and intriguingly, this also revealed little difference in the replication or release of Wt and Δ22 viruses, in a scenario where optimal virus replication requires multiple rounds of replication and spread to other cells in the monolayer (Fig 2A). GFP imaging of cells infected at low multiplicity revealed that the entire monolayer of cells had become GFP positive but without causing CPE, indicating that the Δ22 virus spreads efficiently without affecting the integrity of the cells (Fig 2B). To further visualise the behaviour of the Δ22 virus at low multiplicity and determine if the virus can spread cell-to-cell, HFFF cells were infected with Δ22 (which expresses GFP in place of VP22) or HSV1 GFP-22 at around 20 pfu per well in the presence of 1% human serum to block extracellular virus. Brightfield and GFP fluorescence of representative fields were imaged up to 3 days after infection to investigate virus spread. While HSV1 expressing GFP-22 was seen to spread over time, causing the rounding up of cells as expected (Fig 2C, GFP22), Δ22 failed to cause any obvious CPE (Fig 2C, Δ22, brightfield). Nonetheless, GFP fluorescence was detectable in a cluster of cells at day one, spreading into a much larger area over the next two days (Fig 2C, Δ22, GFP). By contrast, HFFF cells infected with the Δ34 virus at low multiplicity exhibited only individual rounded up cells as late as 3 days after infection, as would be expected as this virus is unable to produce progeny virions (Fig 2D, Δ34). Taken together, these results suggest that despite extreme translational shutoff and no obvious virus-induced pathology, the absence of VP22 has little effect on the propagation of HSV1 in HFFF cells.

**Fig. 2.**
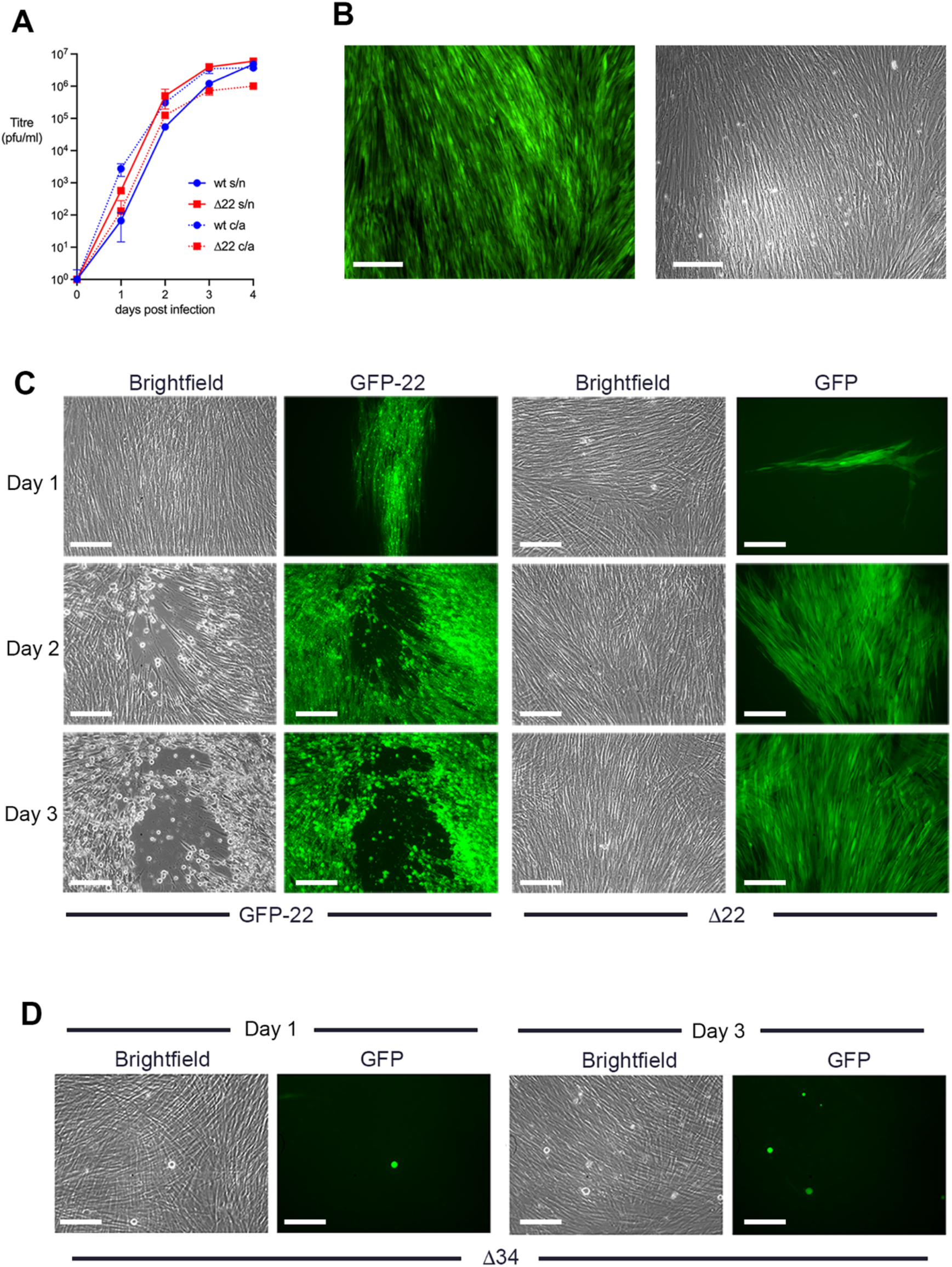
HSV1 spreads in the absence of VP22 without causing CPE. **(A)** HFFF cells were infected with Wt s17 or Δ22 virus at a multiplicity of 0.01, supernatant (s/n) and cell-associated (c/a) virus harvested every day for 4 days and titrated onto Vero cells (mean±SEM, *n* = 3). **(B)** HFFF cells were infected with approximately 20 pfu of HSV1 GFP-Δ22 virus and brightfield and GFP images acquired 3 days later. Scale bar = 100 μm. **(C)** HFFF cells were infected with approximately 20 pfu of HSV1 GFP-22 or Δ22 viruses in the presence of 1% human serum and representative brightfield and GFP images acquired every day for 3 days. Scale bar = 100 μm. **(D)** Confluent HFFF cells were infected with approximately 20 pfu of Δ34 virus and representative brightfield and GFP images acquired at days 1 and 3. Scale bar = 100 μm.

### Point mutations in vhs rescue translational shutoff in Δ22 infected cells

Although the Δ22 virus does not plaque on HFFF cells, plaques spontaneously appear on HFFF cells at a rate of ~ 1 in every 100 pfu, as judged by the original titre on Vero cells [4]. Further analysis of one of these viruses (Δ22*) had previously revealed a single point mutation in the vhs open reading frame (A95T) which had rescued both translation and plaque formation [4]. Taken together with studies from other groups, which have described the rescue of Δ22 replication through spontaneous mutation of vhs [6, 7], and single residue changes in vhs having a profound effect on its activity [17, 18], these results led us to initially hypothesize that the A95T mutation had inactivated the vhs endoribonuclease activity, thereby rescuing late protein synthesis and subsequent virus replication. We have now undertaken a more extensive analysis of this and three additional rescued viruses that were isolated from plaques on HFFF and which formed plaques approaching the size of Wt plaques (Fig 3A).They all express full-length vhs as demonstrated by Western blotting (Fig 3B), indicating that no gross mutations had occurred, but metabolic labelling profiles confirmed that all these viruses had rescued the extreme translational shutoff exhibited by the Δ22 virus, albeit the PP13 virus recovering only slightly from the Δ22 base line (Fig 3C).

**Fig. 3.**
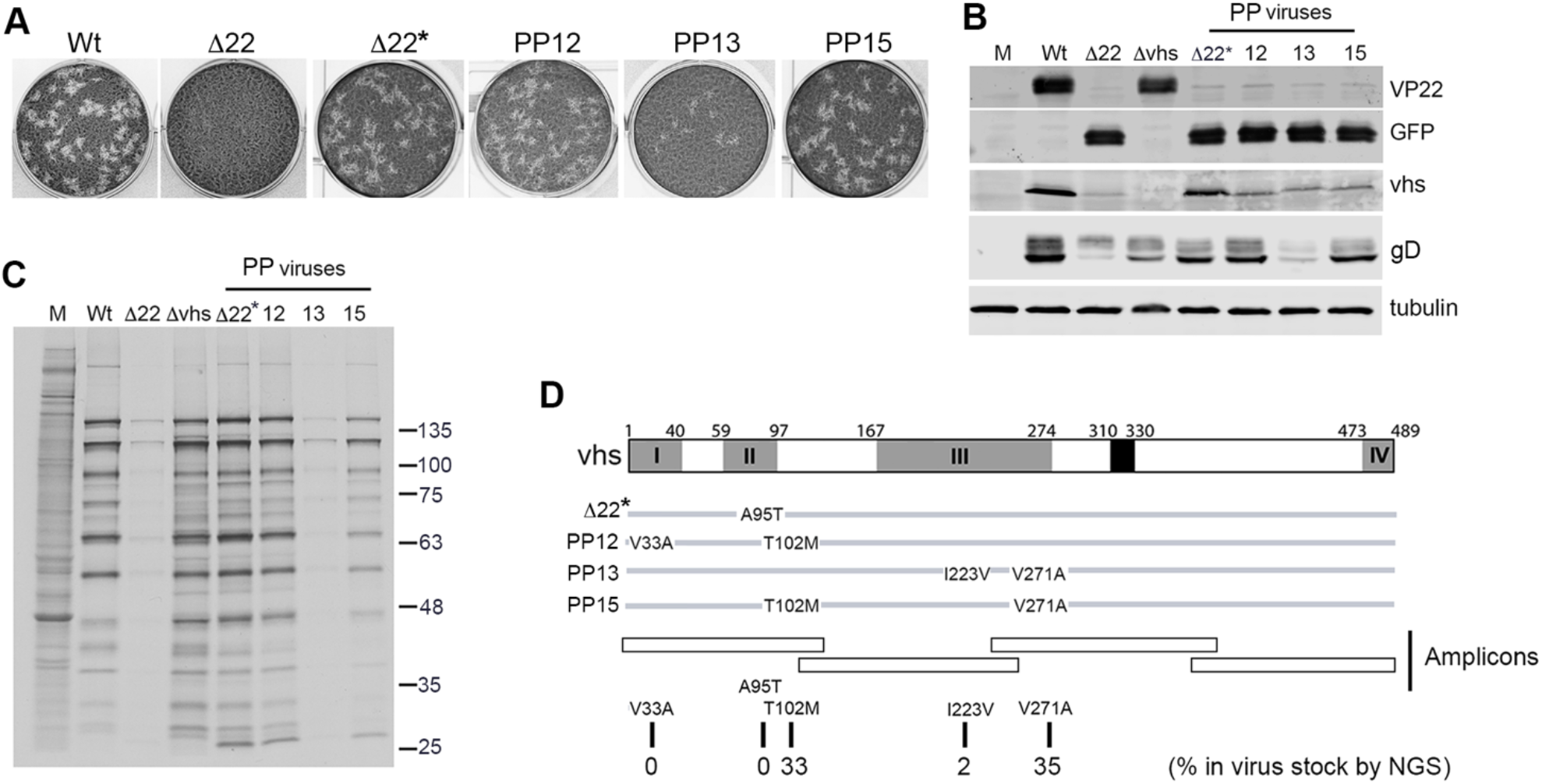
Point mutations in the vhs endoribonuclease rescue plaque formation and translation of Δ22 viruses. **(A)** Virus from four plaques that appeared spontaneously on Δ22-infected HFFF cells was purified and plated onto HFFF cells at around 50 pfu per well (as judged by titre on Vero cells). After 3 days, cells were fixed and stained with crystal violet. **(B)** HFFF cells were infected with the indicated viruses at a multiplicity of 2, harvested at 16 h, subjected to SDS-PAGE and Western blotting for VP22, GFP, vhs and a-tubulin and images acquired with a LICOR Odyssey imaging system. **(C)** HFFF cells were infected with the indicated viruses at a multiplicity of 2, and 16 hours later were incubated in the presence of [35S]-methionine for a further 60 mins. The cells were then lysed and analysed by SDS-PAGE followed by autoradiography. **(D)** A line drawing of the vhs open reading frame indicating the point mutations found in the vhs gene of the rescued Δ22 viruses. The vhs encoding gene, UL41, was amplified by PCR from the submaster stock of our Δ22 virus using four amplicons to cover the entire gene. These amplicons were sequenced by NGS (~40,000 sequences per amplicon) and all variations to the published strain 17 reference sequence (NC001806) scored as the percentage present in the population.

Sequencing of the UL41 gene in these rescue viruses revealed that they all had point-mutations in the vhs open reading frame (Fig 3D). To determine if these variants were present in our original Δ22 virus stock or had arisen during propagation on HFFF cells, we carried out next generation sequencing of four amplicons covering the UL41 gene generated from the genome of our Δ22 virus stock (Fig 3D), revealing that it already contained the T102M and V271A variations at a rate of 33% and 35% respectively, with I223V at a much lower rate of 2% (Fig 3D). No V33A or A95T variations were found by deep sequencing this virus suggesting that they may have arisen spontaneously during propagation on HFFF. Interestingly, direct sequencing of the UL41 gene from 16 viruses isolated from plaques on Vero cells in which this virus is able to plaque, or from nine non-CPE fluorescent foci on HFFF such as those shown in Fig 2C, showed that all viruses contained either the T102M or the V271A variation but none of them contained two mutations. This suggests that each of these single variations in isolation was not sufficient to rescue CPE of this virus in HFFF, and that, with the exception of the A95T variation, a second point mutation was required.

### Restoration of late transcript nuclear export in Δ22 rescue viruses

Given that our previous study had indicated extreme, vhs-dependent, nuclear retention of the virus transcriptome in Δ22 infected cells, we next investigated the subcellular localisation of E (TK) and L (gD) transcripts in HFFF cells infected with the Δ22 rescue viruses by mRNA FISH at 16 hours after infection. As shown previously [4], the IE transcript TK but not the L transcript gD was retained in the nucleus of Wt infected cells, while both transcripts were cytoplasmic in the absence of vhs (Fig 4). By contrast and as shown before [4], both transcripts were almost entirely nuclear in Δ22 infected cells, thereby explaining the observed translational shutoff seen in these cells. In the case of the rescue viruses, virus transcript localisation ranged from both being completely cytoplasmic (Fig 4, Δ22*) similar to that seen in Δvhs infection, to compartmentalisation patterns that were more akin to that of Wt infection than either Δ22 or Δvhs (Fig 4, PP13 and PP15). In all cases there were detectable levels of gD transcripts in the cytoplasm of the rescued virus-infected cells suggesting that the outcome of vhs mutation in the Δ22 virus was the restoration of nuclear export of late transcripts in the absence of VP22, culminating in increased late protein translation.

**Fig. 4.**
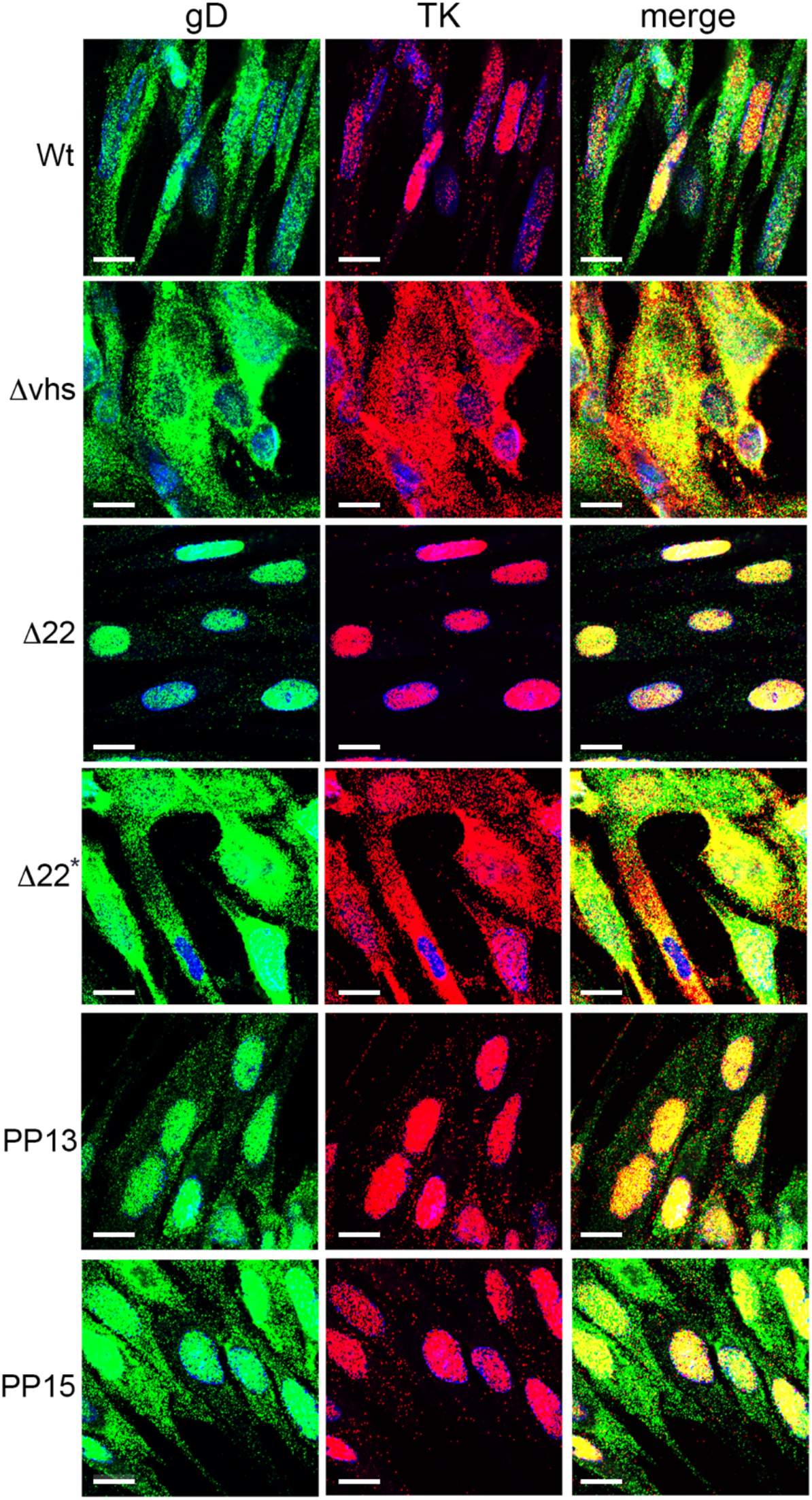
Mutations in vhs rescue viral mRNA export in the absence of VP22. HFFF cells grown in two-well slide chambers were infected with the viruses as indicated at MOI 2 and fixed after 16 hours in 4% paraformaldehyde. Cells were then processed for mRNA FISH using probes specific for E (TK in red) and L (gD in green) transcripts. Nuclei were stained with DAPI (blue). Scale bar= 20μm.

In uninfected cells, PABPC1 has a steady state cytoplasmic localisation, but shuttles between the nucleus and the cytoplasm, binding the polyA tail of mRNAs in the nucleus and being transported out on those tails. It then returns to the nucleus after mRNA turnover in the cytoplasm to be exported again [12]. We have previously shown that in the absence of VP22, the cellular polyA binding protein PABPC1 accumulates to high levels in the nucleus [4, 14], a result we had postulated to be the consequence of the aforementioned nuclear retention of late viral mRNA. We therefore examined the relative compartmentalisation of PABPC1 in HFFF cells infected with Wt, Δvhs, Δ22 or rescue viruses at 10 and 16 hours after infection. As shown before, PABPC1 had partially accumulated in the nucleus of Wt infected cells at 16 h, but remained cytoplasmic in Δvhs infected cells throughout, confirming the role that vhs plays in nuclear relocalisation of PAPBC1 (Fig 5). By contrast, in Δ22 infected cells, PABPC1 had already accumulated in nuclei by 10 h, and was almost entirely nuclear by 16 h, correlating with the extensive accumulation of viral mRNA seen at this time (Fig 5, Δ22). As we have shown before that vhs-induced degradation of mRNA is delayed rather than unrestrained in Δ22 infected HFFF cells [4], this early accumulation of PABPC1 in the nucleus is not a consequence of enhanced degradation of mRNA leading to more PABPC1 entering the nucleus. The relative nuclear accumulation of PABPC1 in cells infected with the Δ22 rescue viruses closely reflected the mRNA nuclear retention seen above: the Δ22* infection was similar to Δvhs, with no nuclear PABPC1; the PP15 virus was similar to Wt, with some PABPC1 in the nucleus; and the two other viruses (PP12 and PP13) caused nuclear accumulation of PABPC1 at levels somewhere between Wt and Δ22 (Fig 5).

**Fig 5.**
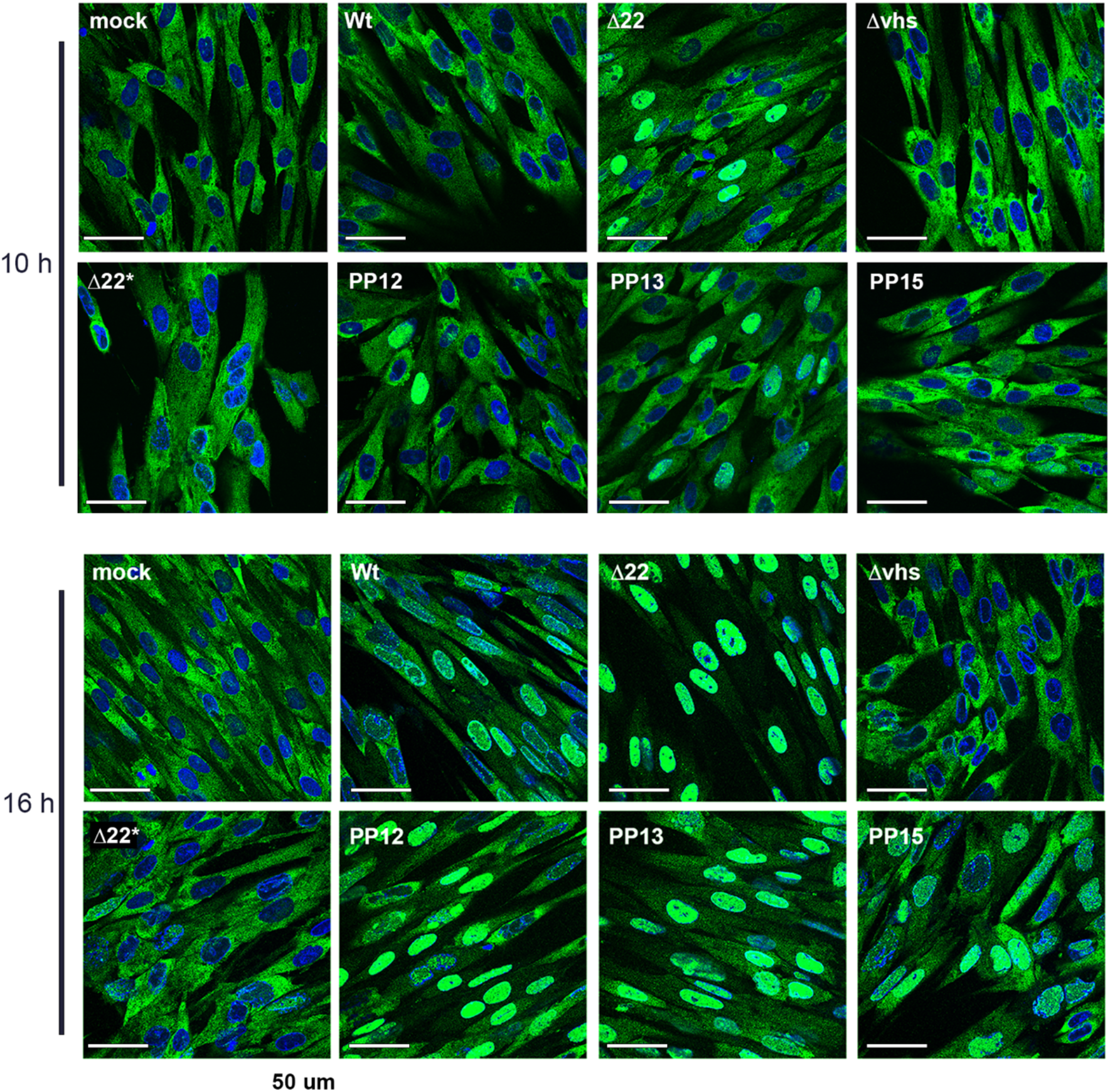
Relative compartmentalisation of PABPC1 in HFFF cells infected with Δ22 rescue viruses. HFFF cells infected with the indicated viruses at MOI 2 were fixed at 10 h and 16 h, stained with an antibody for PABPC1 (green) and nuclei stained with DAPI (blue). Scale bar = 50 μm.

### Rescued Δ22 viruses retain the ability to induce the degradation of cellular mRNA

Our previous work has shown that Δ22 expresses lower levels of L virus transcripts in infected HFFF cells compared to Wt infection, while Δvhs expresses higher levels of IE and E transcripts [4], suggesting that the relative nuclear retention of virus transcripts influences their overall transcription efficiency. To compare the phenotypes of the four rescued viruses, the relative level of a range of virus transcripts across all kinetic classes was compared in HFFF cells infected with Wt, Δ22, Δvhs or each of the four rescued Δ22 viruses. This indicated that all rescued virus infections expressed L transcripts to a level much closer to Wt than Δ22 infection, while Δ22* exhibited high levels of IE and E approaching those found in Δvhs-infected cells (Fig 6A). This suggests that the vhs point mutations in the Δ22 rescue viruses had restored not only the export but also the level of virus transcripts.

**Fig 6.**
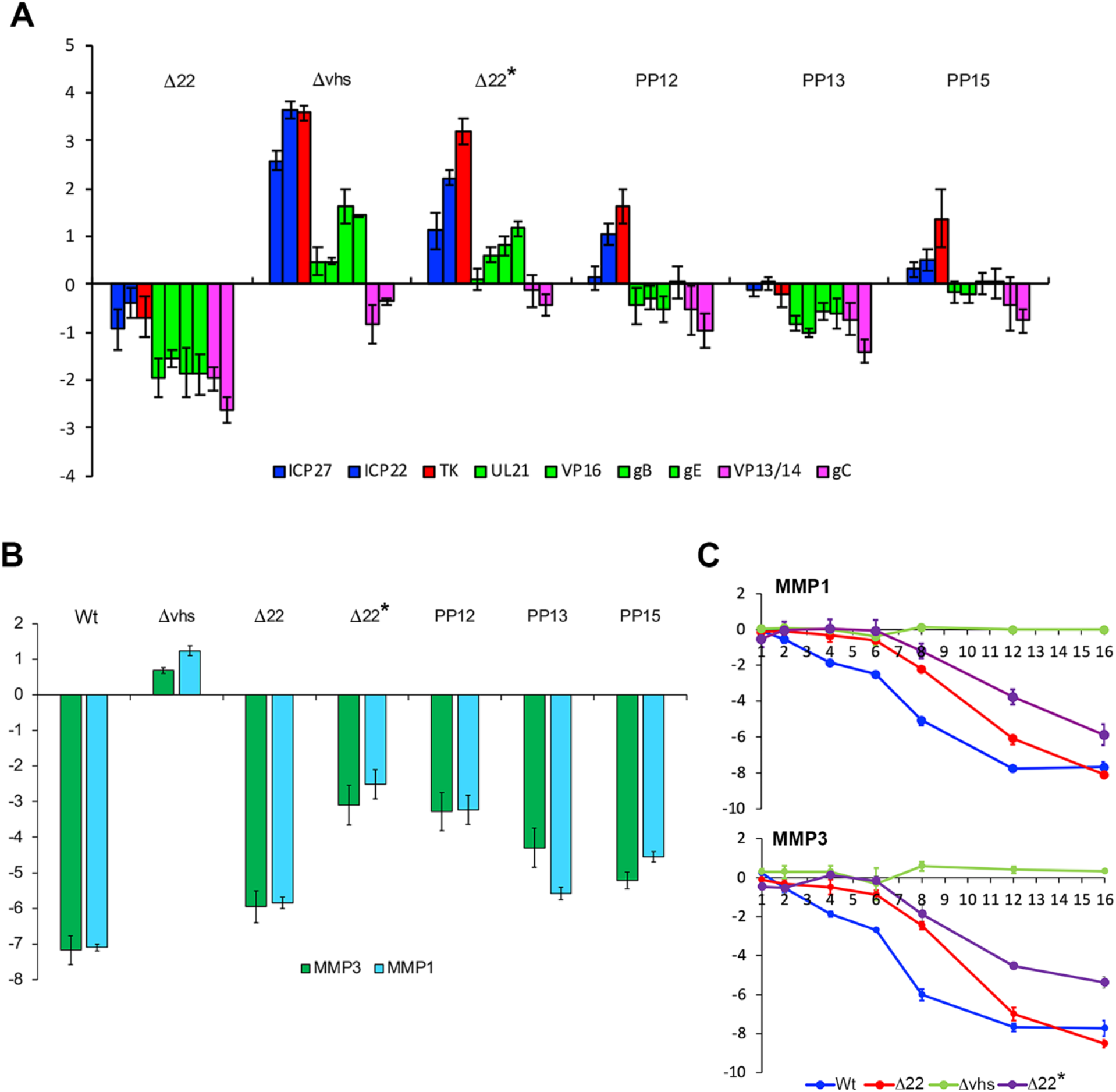
The mutant vhs proteins in rescued Δ22 viruses maintain the ability to degrade cellular mRNA. **(A)** HFFF cells infected with the indicated viruses at a multiplicity of 2 were harvested for total RNA at 16 h and analysed by qRT-PCR for a range of virus transcripts. Results are represented as Log2 fold change to Wt (ΔΔCT), and the mean ± standard error for n = 3 is shown. Transcripts are colour-coded as blue (immediate-early) red (early), green (late) and pink (true late). Part of this data has been presented in a previous publication [4]. **(B)** HFFF cells infected with the indicated viruses at a multiplicity of 2 were harvested for total RNA at 16 h and analysed by qRT-PCR for the cellular MMP1 and MMP3 transcripts. Results are represented as Log2 fold change to uninfected (ΔΔCT), and the mean ± standard error for n = 3 is shown. (**C)** HFFF cells were infected with Wt, Δ22, Δvhs or Δ22* viruses at a multiplicity of 2, and total RNA was harvested at the indicated times (in hours). qRT-PCR was carried out on MMP1 and MMP3 cellular transcripts with relative levels expressed as log2 FC to uninfected (ΔΔCT) over time. The mean and ± standard error for n = 3 is shown. Part of this data has been presented in a previous publication [4].

To determine the relative effect of these additional vhs variants on vhs-induced mRNA degradation, RNA samples were harvested 15 hours after infection and analysed by RT-qPCR for two cellular transcripts we have previously shown to be highly susceptible to vhs degradation - MMP1 and MMP3 [4] (Fig 6B). Surprisingly, all rescue viruses maintained the ability to reduce the levels of both these transcripts compared to Δvhs, with the PP13 and PP15 viruses maintaining activity close to that seen in their parent Δ22 virus (Fig 6B). To look in more detail at the behaviour of the weakest of these vhs variants present in Δ22*, we carried out a time course of relative mRNA levels of MMP1 and MMP3 in comparison to Wt, Δ22 and Δvhs infected HFFF. This confirmed that unlike the situation in Δvhs infected cells where neither the MMP1 nor MMP3 transcript levels were altered during infection, the Δ22 virus caused the gradual decline in these transcripts over time, which as in the Δ22 infection began around 6 hpi and progressed through infection (Fig 6C).

Taking these results together, we have shown that all viruses that were rescued from the Δ22 virus retained vhs activity for mRNA degradation, suggesting that the rescue of Δ22 virus requires vhs mutation to restore late transcript export rather than simple inactivation of vhs endoribonuclease activity. Moreover, this restoration of late transcript export has restored late protein expression together with the ability of these viruses to cause CPE in an environment where virus production had nonetheless not been compromised in the first place.

## Discussion

Primary human fibroblasts offer an excellent model for HSV1 – they are semi-permissive to infection, having the capacity to restrict HSV1 at early stages, and they have a functioning interferon pathway [19]. They also exhibit extreme CPE in response to HSV1 infection with profound changes to cell architecture from long spindle-like to small rounded up cells (Fig 1E). It was therefore intriguing to discover that our mutant Δ22 HSV1 virus was able to enter, replicate and spread within HFFF cells in a similar fashion to Wt virus but without causing CPE. CPE is generally considered to be a combination of gross morphological changes including cytoskeletal and membrane reorganisation, together with physiological and biochemical changes caused by the virus hijacking cellular activities. The absence of CPE and plaque formation in Δ22 infected HFFF cells, which we had originally assumed to indicate attenuation, correlates with late translational shutoff suggesting that CPE is caused by the high levels of late virus proteins, or specific proteins, made within the cell. Indeed, an HSV1 deleted for ICP34.5 – a protein whose absence results in translational shutoff via the PKR-eiF2α pathway – also fails to cause CPE in human fibroblasts [20]. Moreover, it has recently been reported that HSV1 which fails to cause CPE in culture has been isolated from patients on acyclovir therapy [21], and despite expressing very low levels of all virus proteins, these viruses can be propagated in culture. One important outstanding question from all of these CPE-negative viruses is how newly assembled virions transfer from cell-to-cell to propagate infection in the absence of cellular alterations, raising the possibility that virus egress and release from infected cells does not require any specific injury to the cells.

Despite efficient replication of the Δ22 virus in the absence of CPE, there appears to be a clear pressure on the virus to regain a high level of late protein translation. Unlike infection of a Δ34.5 virus, translational shutoff in the absence of VP22 does not appear to involve PKR restriction of the translation machinery but correlates with the entrapment of the virus transcriptome in the nucleus of infected cells [4]. The spontaneous appearance of CPE-causing virus in the Δ22 virus seems to be a consequence of SNPs in the vhs open reading frame. These SNPs cluster within the N-terminal half of vhs (Fig 7), fitting with other mutations found in previous Δ22 virus studies which cluster in conserved domain III of the protein [18, 22, 23]. These have been interpreted previously as mutations that inactivate vhs endoribonuclease activity [6, 7, 24] and in support of this, analysis of vhs sequences from 26 published strains of HSV1 showed that naturally occurring SNPs cluster within the C-terminus of the protein while the N-terminus is highly conserved (Fig 7). Moreover, in the course of this study, the vhs gene from ten clinical isolates of HSV1 [25] were also sequenced, with seven of them identical to strain 17, and three containing SNPs in the C-terminal half already identified in the published sequences. Nonetheless, our rescue viruses retain the ability to degrade cellular mRNA – a readout for endoribonuclease activity – whilst regaining the ability to export late mRNA to the cytoplasm for access to the translation machinery. As such, these mutants have separated the contribution that vhs makes to nuclear export and mRNA degradation. This is further emphasized by the early relocalisation of PABPC1 to the nucleus of Δ22 infected cells at a time when mRNA degradation is reduced compared to Wt infection (Fig 6B) [4], suggesting that mRNA and hence PABPC1 retention in the nucleus can be uncoupled from mRNA degradation. Moreover, we have also recently described a virus expressing vhs tagged with GFP, that fails to degrade cellular mRNA, is unable to relocalise PABPC1 during infection, but can still cause nuclear retention of IE and E transcripts [26].

**Fig 7.**
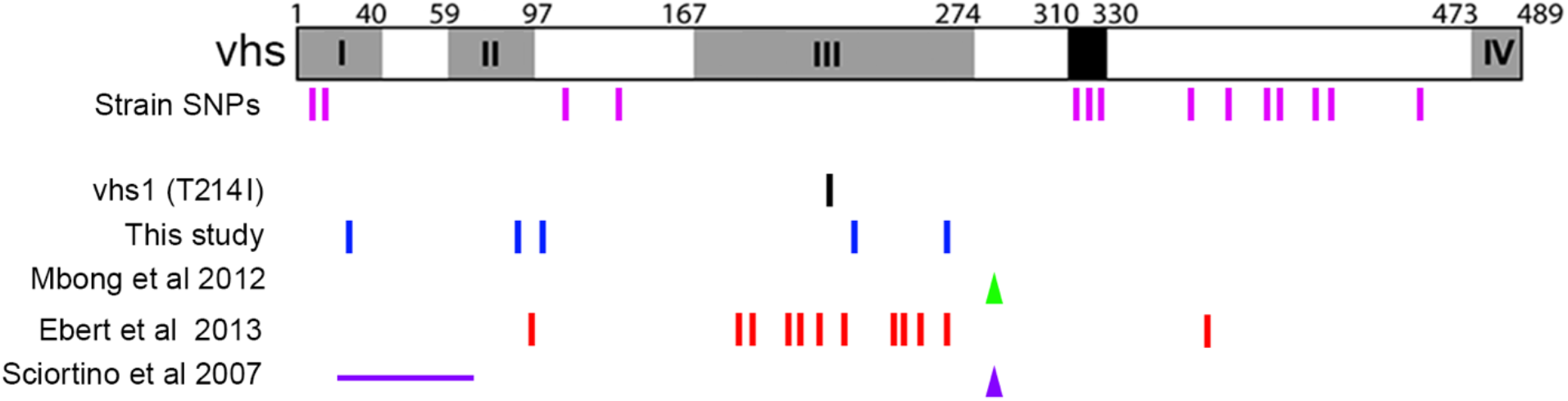
Summary of SNPs that have been found previously in the vhs open reading frame: pink – SNPs found in 12 out of 26 published HSV1 sequences compared to strain 17(Table 3); black - the well-characterised vhs1 mutation (T214I) that has been found to abrogate vhs activity [17]; blue - SNPs found in our Δ22 virus in this study; green – frameshift found in Δ22 virus rescued from a strain F BAC [7]; red – SNPs found in a strain 17 BAC-constructed Δ22 virus after transfection and rescue of multiple viruses [24]; purple - deletion and frameshift found in strain F Δ22 virus rescued from a BAC [6]. Vertical line – SNP. Triangle – frame shift. Horizontal line – deletion.

Although we do not yet understand the molecular basis of these complex phenotypes, the broad range of SNPs across the N-terminus of vhs will allow us to further separate the activities of mRNA degradation, PABPC1 relocalisation and mRNA nuclear retention. In addition, it is important to note that although our Δ22 virus which is based on strain 17 does not plaque in human fibroblasts, it is able to form small plaques, and hence CPE, on Vero cells [4, 27]. By contrast, other reports of HSV1 VP22 deletion viruses based on strain F have indicated that these viruses are unable to plaque even on Vero cells [5, 6]. The VP22 and vhs sequences in these two strains have several SNPs compared to strain 17, providing scope for strain variation in the interplay between these proteins and the machinery/pathways involved in regulating protein translation.

The work presented here adds further weight to the important roles that vhs and VP22 play in the co-ordinated regulation of mRNA localisation and translation. HSV1 also expresses the IE protein ICP27 which is known to be essential for late protein expression [28], and is required for mRNA export from the nucleus by binding TAP/NXE1 to engage with the cellular Aly/REF export pathway [29–31]. It is therefore paradoxical that one activity of vhs is to retain virus transcripts in the nucleus, suggesting that vhs and ICP27 may work in opposition to each other. Going forward, it will now be important to establish how vhs (and VP22) intersect with the nuclear export activity of ICP27 to co-ordinate virus gene expression via mRNA compartmentalisation.

The question remains as to why there is a selective pressure on the virus to restore late translation/CPE through vhs mutation in the absence of VP22, if the virus is able to replicate and spread as efficiently as Wt virus. One could imagine that a virus that propagates itself without causing damage to its host cell might be able to survive “under the radar” of host sensing mechanisms and antiviral measures. However, the rapid and reproducible appearance of secondary mutations within vhs suggests that the production of a large amount of CPE-causing structural proteins is advantageous to the virus, over and above the requirement for virus assembly. It is therefore likely that structural proteins or a subset of them are required to maintain a favourable environment for virus survival, and as suggested elsewhere may reflect the ongoing battle between host and virus during virus infection [32]. As such, the unexpected results described here may ultimately prove highly informative about the range of mechanisms by which HSV1 overcomes host defences.

## Methods

### Cells and Viruses

HFFF and Vero cells (both obtained from European Collection of Authenticated Cell Cultures - ECACC) were cultured in DMEM supplemented with 10% foetal bovine serum (Invitrogen). Viruses were routinely propagated in Vero cells, with titrations carried out in DMEM supplemented with 2% foetal bovine serum and 1% human serum. HSV1 strain 17 (s17) was used routinely. The s17 derived VP22 deletion mutant (Δ22) and the vhs knockout virus (Δvhs) have been described before [27, 33]. The four Δ22 rescue viruses (Δ22*, PP12, PP13 and PP15) were isolated from plaques that appeared spontaneously on HFFF cells.

### Antibodies & reagents

Our VP22 (AGV031) antibody has been described elsewhere [34, 35]. Other antibodies were kindly provided as follows: gD (LP14), VP16 (LP1) and gM, Tony Minson and Colin Crump (University of Cambridge, UK); vhs, Duncan Wilson (Albert Einstein College of Medicine, USA); gE, David Johnson (Oregon Health and Science University, Portland, USA). Other antibodies were purchased commercially: a-tubulin (Sigma), GFP (Clontech), PABPC1 and ICP4 (Santa Cruz), and gC (Abcam). Horseradish peroxidase-conjugated secondary antibodies were from Bio-Rad Laboratories and IRDye secondary antibodies were from LICOR Biosciences.

### SDS-PAGE and Western blotting

Protein samples were analysed by SDS-polyacrylamide gel electrophoresis and transferred to nitrocellulose membrane for Western blot analysis. Western blots were developed using SuperSignal West Pico chemiluminescent substrate followed by exposure to X-ray film, or by imaging on a LICOR Odyssey Imaging system.

### Metabolic labelling of infected cells

Cells grown in 3cm dishes were infected at a multiplicity of 2, and at indicated times were washed and incubated for 30 mins in methionine-free DMEM before adding 50μCi of L-[35S]-methionine (Perkin Elmer) for a further 30 min. Cells were then washed in PBS and total lysates analysed by SDS-polyacrylamide gel electrophoresis. Following fixation in 50% v/v ethanol and 10% v/v acetic acid, the gel was vacuum dried onto Whatman filter paper and exposed to X-ray film overnight.

### Quantitative RT-PCR (RT-qPCR)

Total RNA was extracted from cells using Qiagen RNeasy kit. Excess DNA was removed by incubation with DNase I (Invitrogen) for 15 min at room temperature, followed by inactivation for 10 min at 65°C in 25 nM of EDTA. Superscript III (Invitrogen) was used to synthesise cDNA using random primers according to manufacturer’s instructions. All qRT-PCR assays were carried out in 96-well plates using MESA Blue qPCR MasterMix Plus for SYBR Assay (Eurogentec). Primers for viral genes are shown in Table 1. Primers for cellular genes MMP1 and MMP3 were obtained from SinoBiological (HP100549 and HP100493 respectively). Cycling was carried out in a Lightcycler (Roche), and relative expression was determined using the ΔΔCT method [36], using 18s RNA as reference.

**Table 1.**
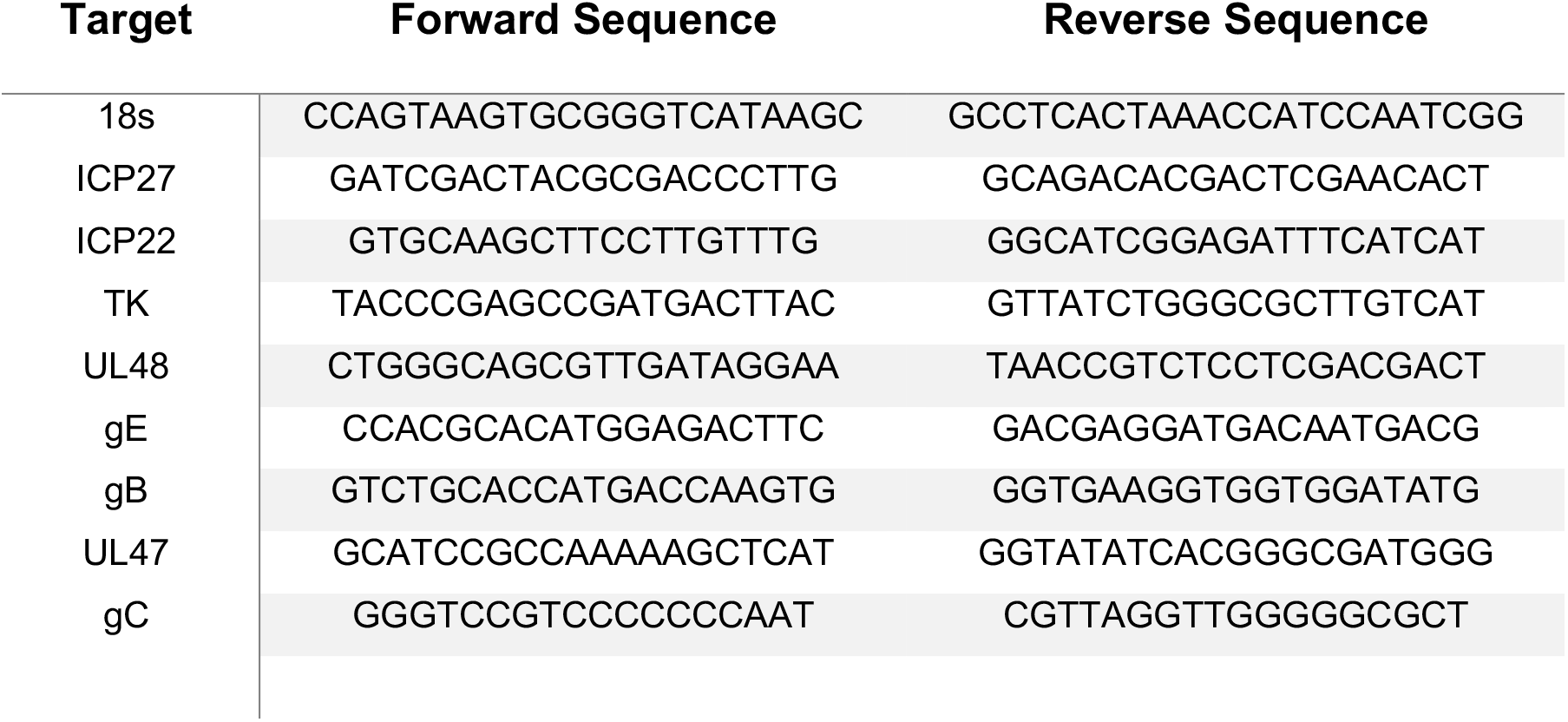
Primer sequences for RT-qPCR of indicated virus genes

### Quantification of viral DNA

HFFF cells infected at MOI 3 were acid washed 1 h after infection to inactivate unpenetrated virus, then harvested at 2 or 16 h after infection. DNA was harvested using the DNeasy blood and tissue kit (Qiagen), and qPCR assays were carried out in a LightCycler96 system (Roche), using MESA BLUE qPCR kit for SYBR assay (Eurogentec) according to the manufacturer’s instructions with primers for 18S (see Table 1) and HSV1 *UL48* gene.

### Amplicon Sequencing

The full-length *UL41* gene was amplified in four fragments by PCR from our virus submaster stock of the Δ22 virus, using the primers shown in Table 2. Each PCR fragment was purified and subjected to amplicon next generation sequencing (Genewiz) providing around 40,000 reads per amplicon.

**Table 2.**
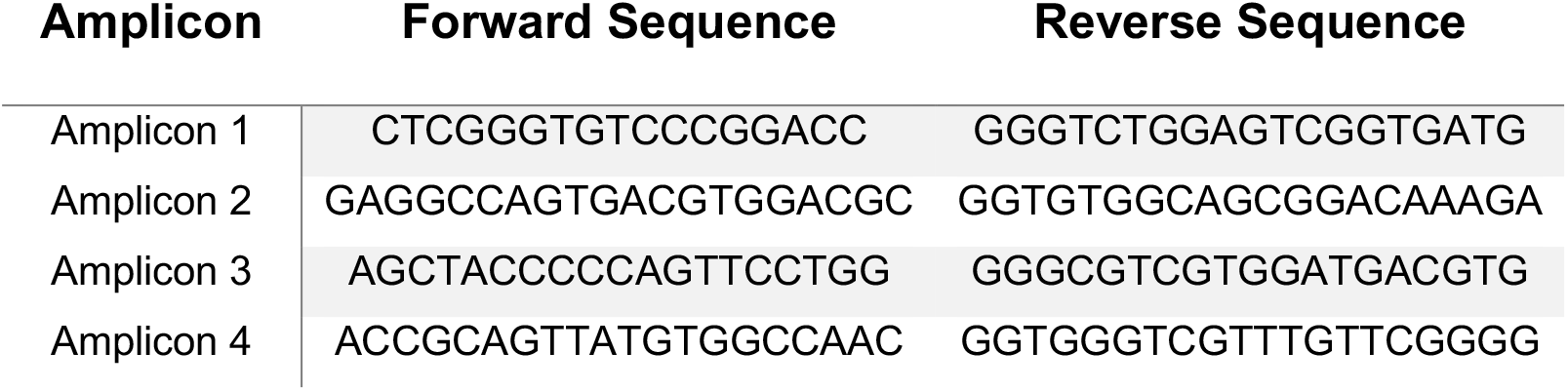
Primers used to amplify four overlapping fragments of the vhs open reading frame, UL41

### Immunofluorescence

Cells for immunofluorescence were grown on coverslips and fixed with 4% paraformaldehyde in PBS for 20 min at room temperature, followed by permeabilisation with 0.5% Triton-X100 for 10 min. Fixed cells were blocked by incubation in PBS with 10% newborn calf serum for 20 min, before the addition of primary antibody in PBS with 10% serum, and a further 30-min incubation. After extensive washing with PBS, the appropriate Alexafluor conjugated secondary antibody was added in PBS with 10% serum and incubated for a further 15 min. The coverslips were washed extensively in PBS and mounted in Mowiol containing DAPI to stain nuclei. Images were acquired using a Nikon A1 confocal microscope and processed using ImageJ software [37].

### Fluorescent *in situ* hybridisation (FISH) of mRNA

Cells were grown in 2-well slide chambers (Fisher Scientific) and infected with virus. At the appropriate time, cells were fixed for 20 min in 4% PFA, then dehydrated by sequential 5 min incubations in 50%, 70% and 100% ethanol. FISH was then carried out using Applied Cell Diagnostics (ACD) RNAscope reagents according to manufacturer’s instructions. Briefly, cells were rehydrated by sequential 2 min incubations in 70%, 50% ethanol and PBS, and treated for 30 min at 37 °C with DNase, followed by 15 min at room temperature with protease. Cells were then incubated for 2 h at 40 °C with RNAscope probes for ICP0, TK or glycoprotein D as designed by Advanced Cell Diagnostics, ACD, followed by washes and amplification stages according to instructions. After incubation with the final fluorescent probe, the cells were mounted in Mowiol containing DAPI to stain nuclei, and images acquired with a Nikon A2 inverted confocal microscope and processed using Adobe Photoshop software.

### Microscopy

Images were acquired on a Nikon A2 confocal microscope or CCD camera system on an inverted Zeiss TV100 microscope and processed using Image J and Adobe Photoshop software.

### UL41 sequence comparison

The UL41 gene sequence from 26 previously published HSV1 isolates (Table 3; [38–41]) were aligned to the sequence from strain 17 and single nucleotide polymorphisms (SNPs) in comparison to strain 17 vhs were identified. In the course of this study, UL41 from ten clinical isolates of HSV1 [25] was also sequenced. Seven of these were identical to strain 17 UL41, while three contained SNPs identified in the C-terminal domain of previously sequenced HSV1 genomes denoted above.

**Table 3.** Published HSV-1 sequences used in this study for comparison of vhs sequences.

| Strain | Accession Number | Strain | Accession Number |
| --- | --- | --- | --- |
| s17 | NC001806 | TFT401 | JN420337 |
| Sc16 | KX946970 | CJ394 | JN420340 |
| F | KM222725 | McKrae | JX142173 |
| H129 | GU734772 | KOS63 | KT425110 |
| E06 | HM585496 | KOS79 | KT425109 |
| E10 | HM585499 | CM1 | KX791792 |
| E11 | HM585500 | HSZP | Z72337 |
| E15 | HM585503 | S23 | HM585512 |
| E22 | HM585504 | S25 | HM585513 |
| E23 | HM585505 | R11 | HM585514 |
| E25 | HM585506 | R62 | HM585515 |
| E35 | HM585507 | CR38 | HM585508 |
| 134 | JN400093 | HF10 | DQ889502 |

## Acknowledgments

The authors thank Duncan Wilson, David Johnson, Tony Minson and Colin Crump for antibodies used in this study. This work was supported by grants from the Medical Research Council, UK (MR/M011607/1 & MR/T0001038/1).

